# Large-scale forward genetics reveals genetically separable components of plant cell proliferative capacity

**DOI:** 10.64898/2026.10.30.762808

**Authors:** Fuensanta Verdú-Navarro, Ángela Espín-Veracruz, Julia Weiss, Joanna Kacprzyk, Pedro J. Navarro-Lorente, Dolores Parras-Burgos, David Borraz-Moron, Eduardo Salas-Fernández, Marcos Egea-Cortines

**Affiliations:** Universidad Politécnica de Cartagena, member of European University of Technology EUT+, Instituto de Biotecnología Vegetal, 30203 Cartagena, Spain; Bioprocess Pilot Plant, BIONET, 30320 Fuente Álamo, Spain; School of Biology and Environmental Science, University College Dublin, D04 VIW4, Ireland; Universidad Politécnica de Cartagena, member of European University of Technology EUT+, División de Sistemas e Ingeniería Electrónica, 30203 Cartagena, Spain; Universidad Politécnica de Cartagena, member of European University of Technology EUT+, Grupo de Crear y Diseñar en Ingeniería, 30203 Cartagena, Spain

**Keywords:** Callus proliferation, Forward genetics, High-throughput phenotyping, Computer vision, T-DNA insertion mutants, Enhanced biomass accumulation, Plant cell suspension culture, Sustained proliferative capacity, Serial subculture, Plant cell bioprocessing

## Abstract

Plant cell cultures underpin regeneration, transformation and emerging applications in biotechnology and plant cell agriculture, yet the genetic determinants of proliferative performance remain poorly characterized. Here, we combined large-scale forward genetics with automated image-based phenotyping to quantify 27,648 calli representing 2,880 Arabidopsis thaliana T-DNA insertion lines. Of these, 2,456 lines produced calli smaller than the WT population mean, whereas only 190 produced larger calli, revealing a strongly asymmetric phenotypic distribution. Enhanced callus growth was not consistently associated with increased vegetative growth, indicating that these traits can be genetically uncoupled. Validation of selected high-growth mutants in suspension culture identified lines with enhanced biomass accumulation, while serial propagation revealed independent variation in sustained proliferative capacity. Thus, callus growth, whole-plant growth, short-term suspension biomass accumulation and sustained proliferation represent distinct, not necessarily coupled components of proliferative performance. These findings establish a scalable framework for identifying genetic determinants relevant to the development of improved plant cell production lines.

**Highlights:**

- Established a high-throughput platform for quantitative phenotyping of plant callus growth.
- Screened 2,880 Arabidopsis T-DNA lines, revealing extensive genetic variation in callus proliferative capacity.
- Identified mutants with enhanced biomass accumulation or sustained proliferative capacity in suspension culture.
- Established a framework for the genetic improvement of plant cell production platforms.

Graphical Abstract Overwiew of the study. Image created with Biorender.

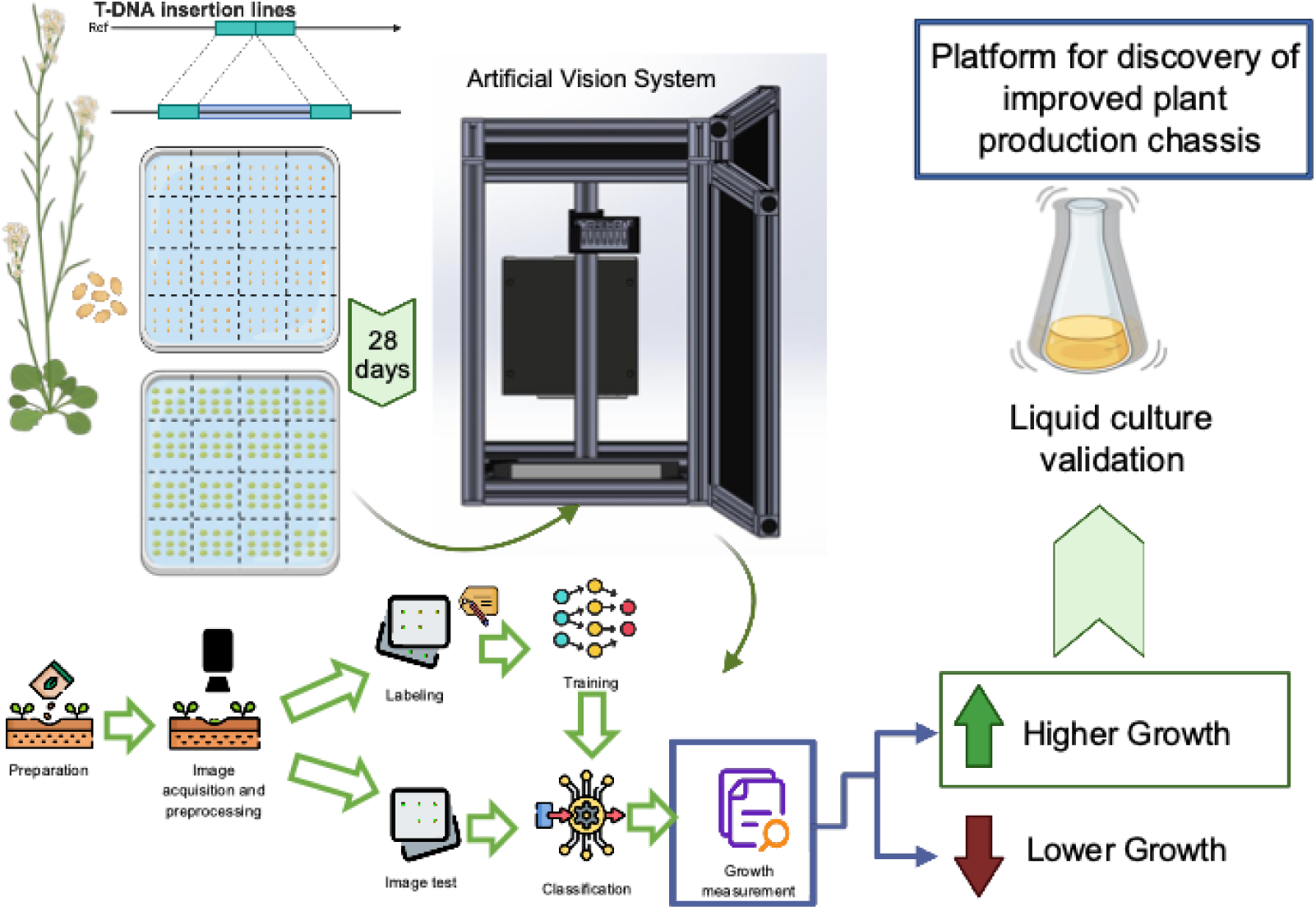

## Introduction

Plant cell and callus cultures underpin plant transformation, regeneration and an expanding range of industrial applications, including the production of recombinant proteins, high-value metabolites and emerging cellular agriculture products ^1–3^. Despite their growing industrial relevance, efforts to improve plant cell cultures have focused predominantly on medium optimization, elicitation strategies and bioprocess engineering. Unlike microbial and mammalian production hosts, plant cell cultures have not been subjected to systematic large-scale genetic improvement for enhanced proliferative capacity.

Genetic studies of plant cell cultures have focused predominantly on regeneration competence rather than industrial performance. Landmark discoveries identifying regulators such as PLT, WOX and WIND1 established the molecular basis of callus induction, totipotency and de novo shoot regeneration, enabling regeneration of previously recalcitrant species ^4,5^. Likewise, manipulation of individual cell-cycle regulators demonstrated that callus proliferation can be genetically modified. For example, overexpression of *CYCD3* enhances callus induction and growth in Arabidopsis ^6,7^., while expression of a rice *R2 CDK*-activating kinase is sufficient to induce callus formation in tobacco explants in the absence of cytokinin ^8^. These studies established that proliferation is genetically tuneable, but they examined individual developmental regulators rather than the genetic architecture of proliferative capacity itself. Consequently, proliferative capacity has not been systematically explored as a quantitative industrial trait for improving transformation efficiency, biomass accumulation and the performance of plant cell production platforms.

A major barrier to treating proliferative capacity as a quantitative trait has been the lack of scalable phenotyping protocols. Unlike traits such as flowering time or plant architecture, callus growth has traditionally required manual image analysis or destructive measurements, limiting the number of genotypes that could be evaluated. Recent advances in automated imaging and AI-based image segmentation now enable objective, high-throughput quantification of callus size in large image datasets ^9–11^. Combined with the availability of genome-wide Arabidopsis T-DNA insertion collections ^12^, these technologies now make forward genetic dissection of proliferative capacity feasible at population scale.

Here, we combined large-scale forward genetics in *Arabidopsis thaliana* with standardized callus induction and deep-learning-based object detection and image segmentation to systematically identify genetic determinants of callus growth. By screening 2,880 homozygous T-DNA insertion lines, we asked whether growth-related phenotypes identified in a high-throughput image-based screen could reveal candidate loci for improving plant cell cultures. We further evaluated representative high-growth mutants to determine whether increased callus size translated into enhanced biomass accumulation and sustained proliferation in suspension culture. Together, our results establish a scalable framework for identifying genetic determinants of plant cell proliferative performance and candidate targets for the development of improved plant cell production lines.

## Results

### Testing cell cycle related genes as a proof of concept

Previous studies have demonstrated that mutations in *Arabidopsis thaliana KIP RELATED PROTEIN KRP* (*INHIBITOR OF CYCLIN KINASE-ICK*) genes, which encode cyclin-dependent kinase inhibitors, result in enhanced cell proliferation and increased organ size. Double and triple *KRP* mutants display enlarged leaves ^13^. Furthermore quintuple ICK/KRP knockouts exhibit increased cell proliferation, larger organs and seeds, and enhanced callus growth ^14,15^.

Based on this evidence, we hypothesized that T-DNA insertion lines affecting cell-cycle-related genes would display detectable alterations in callus size under our induction conditions. Because the aim was to develop a scalable forward screen, calli were induced directly from seeds ^16,17^, avoiding the need to generate leaf explants for each mutant line and biological replicate. The callus induction medium contained Murashige Skoog salts (MS), B5 vitamins 2,4 Dichloro phenoxyacetic acid and 6-Benzyl Aminopurine. To test this as a proof of concept, we selected 13 homozygous T-DNA insertion lines targeting cell cycle genes including *AtKRP1*, *AtKRP5*, *AtKRP7*, and *AtSIAMESE* (Table S1). Mutant and wild-type seeds were randomly distributed across plates (Figure S1), and germination rate and callus growth were evaluated by visual inspection.

Several mutants displayed altered germination and readily detectable differences in callus size relative to Col-0 (Figure S1). Mutations in *SIM*, *AtKRP1*, and one *AtKRP7* allele produced smaller calli, whereas a second independent *AtKRP7* allele exhibited increased callus size. Previous studies reported enhanced callus development only in higher-order *KRP* mutants ^15^, whereas our results indicate that individual alleles can generate detectable phenotypes under the screening conditions employed here. These observations validated the experimental design and indicated that a genome-scale forward genetic screen would be feasible despite the functional redundancy expected among cell-cycle regulators.

### A scalable imaging and AI pipeline enables high-throughput callus phenotyping

Reliable quantitative comparison of callus size across thousands of genotypes requires highly standardized image acquisition. Variations in illumination, camera position or plate orientation introduce systematic errors that can obscure relatively small phenotypic differences. To minimize these sources of variation, we developed a custom-built dark chamber housing the imaging system (Figure 1 a,b). Plates were illuminated using uniform white backlight illumination and imaged with an OAK-1 RGB camera positioned perpendicular to the plate under fixed geometric conditions, enabling consistent pixel-based area quantification.

**Figure 1.**
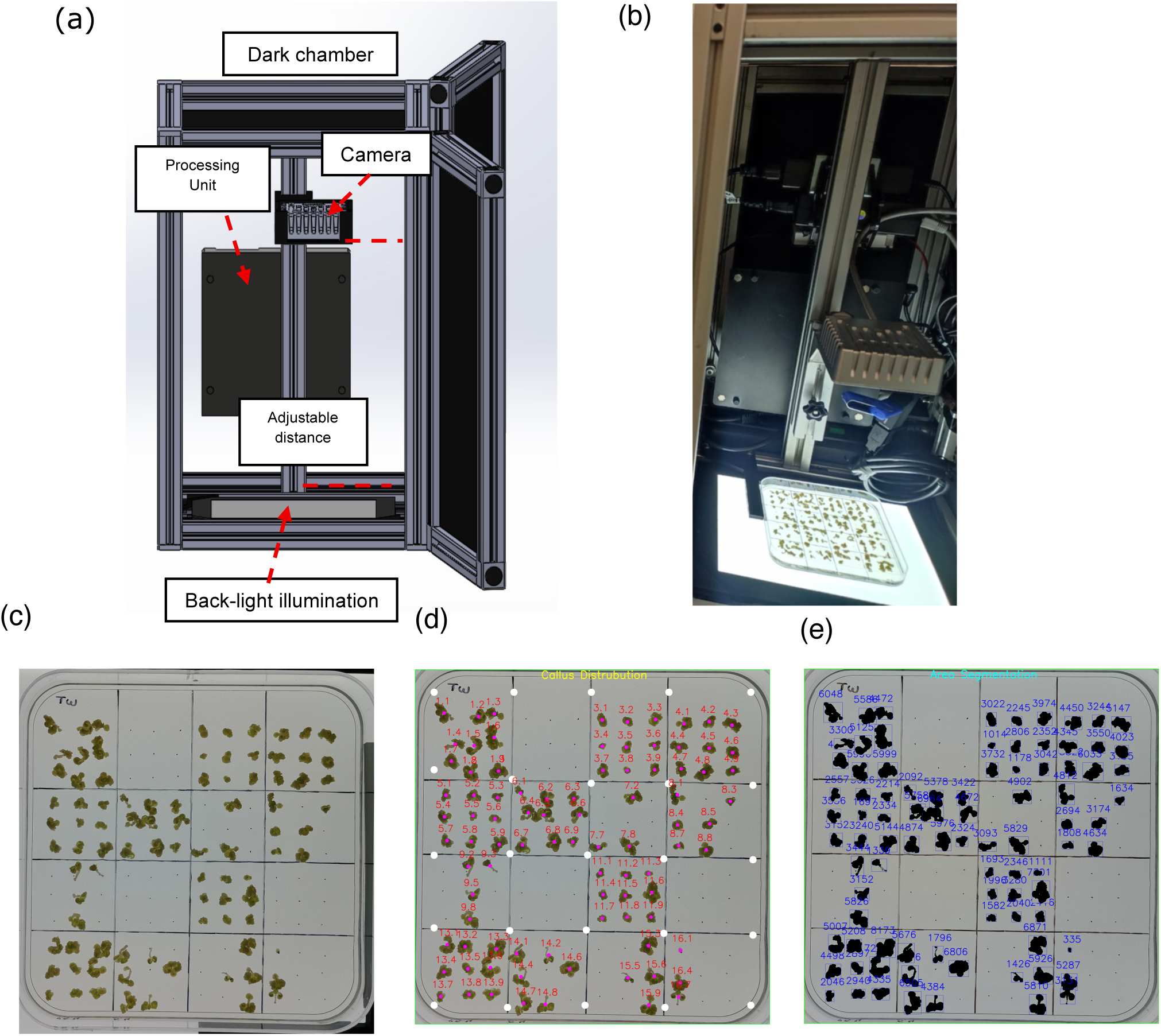
Automated image acquisition and quantitative phenotyping of callus growth. (a) Schematic representation of the image acquisition device. (b) Photograph of the imaging apparatus. (c) Image of a screening plate. (d) Detection of individual calli using the trained YOLO11 model. Red labels indicate automatically assigned sector and callus identifiers. (e) Segmentation and area quantification of individual calli. Blue labels indicate the calculated area of each callus in pixels.

Captured images were transmitted to an NVIDIA Jetson TX2 processing unit, where preprocessing, image acquisition control, and inference were executed through a custom graphical interface (see Materials and Methods).

To enable quantitative comparison across thousands of independent genotypes, key experimental variables were standardized, including callus induction conditions, seed distribution, plate orientation during image acquisition and automated genotype assignment.

Prior to large-scale screening, a deep-learning model was trained to distinguish proliferating calli from non-germinated seeds. A ground-truth dataset comprising 1,230 manually annotated calli, together with associated non-germinated-seed annotations from 26 Petri dish images, was used for model training and evaluation. The resulting model accurately detected calli and non-germinated seeds across all plates (Figure 1c–e). Following object detection, individual calli were segmented using HSV color-space thresholding and contour extraction, enabling pixel-based area quantification.

Detected calli were automatically assigned to their corresponding genotypes based on their position within the predefined plate sectors. Callus area measurements were exported for downstream statistical analysis, enabling standardized high-throughput quantification of callus size across the mutant population.

### High-throughput screening of a T-DNA insertion population for callus proliferation

The screening platform was applied to 2,880 homozygous Arabidopsis T-DNA insertion lines distributed across 192 Petri plates (12 × 12 cm) (Table S2), each containing 15 mutant lines and one plate-matched Col-0 control.

Of the 2,880 lines screened, 723 showed reduced or null germination (Table S3). After excluding 234 lines with complete absence of germination, a total of 27,648 calli were analyzed 28 days after plating, comprising 1,728 WT and 25,920 mutant calli (Figure 2 a). Across the full dataset, the distribution of callus areas in the mutagenized population was shifted toward smaller values relative to WT (*t*-test, *P* < 2.2 × 10⁻¹⁶; Figure 2a). The screen identified mutants displaying reduced germination as well as both reduced and increased callus size (Figure 2b–d).

**Figure 2.**
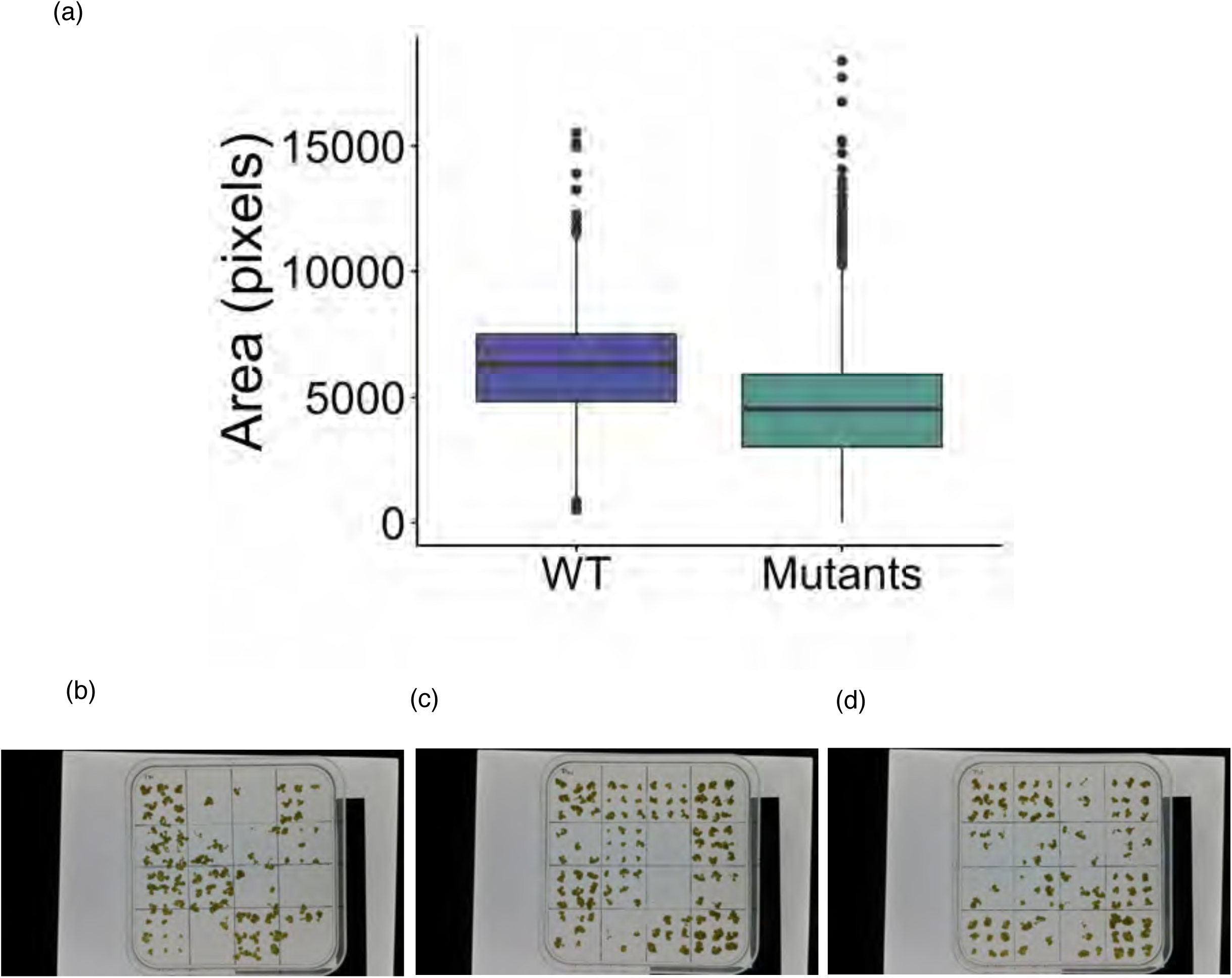
Large-scale screening reveals extensive genetic variation in callus growth. (a) Distribution of callus area (pixels) in 1,728 wild-type (Col-0) and 25,920 mutant calli. The distributions differed significantly (two-sample t-test, *P* < 2.2 × 10⁻¹⁶). Representative screening plates illustrating (b) lines with low germination rate, (c) a mutant line producing smaller calli than Col-0, and (d) a mutant line producing larger calli than Col-0. In each plate, the Col-0 control occupies the upper-left sector.

Lines with fewer than four germinated seedlings were retained for descriptive analysis but excluded from quantitative size comparisons. Thirteen plates in which the WT control displayed abnormal growth were identified as batch outliers and excluded from subsequent analyses.

Using the mean WT callus area (6,430 pixels) as a population reference, 2,456 mutant lines produced smaller calli than WT, whereas only 190 produced larger calli (Tables S4 and S5), indicating a strong asymmetry in the distribution of growth phenotypes. To identify candidate lines while accounting for plate-to-plate variation, each mutant was subsequently compared with the Col-0 control grown on the same plate using two-sided *t*-tests. At a nominal significance threshold of *P* < 0.05, 988 lines showed reduced callus growth and 19 showed increased callus growth relative to their plate-matched WT controls (Table S6).

After Benjamini–Hochberg correction for multiple testing, one high-growth line remained significant at FDR < 0.05. Because the primary screen was designed to identify candidates for subsequent experimental validation, high-growth lines were prioritized on the basis of their plate-matched phenotype and confirmation of the corresponding T-DNA insertion. Fifteen high-growth lines were ultimately advanced for further characterization (Table S7; Figure 3).

**Figure 3.**
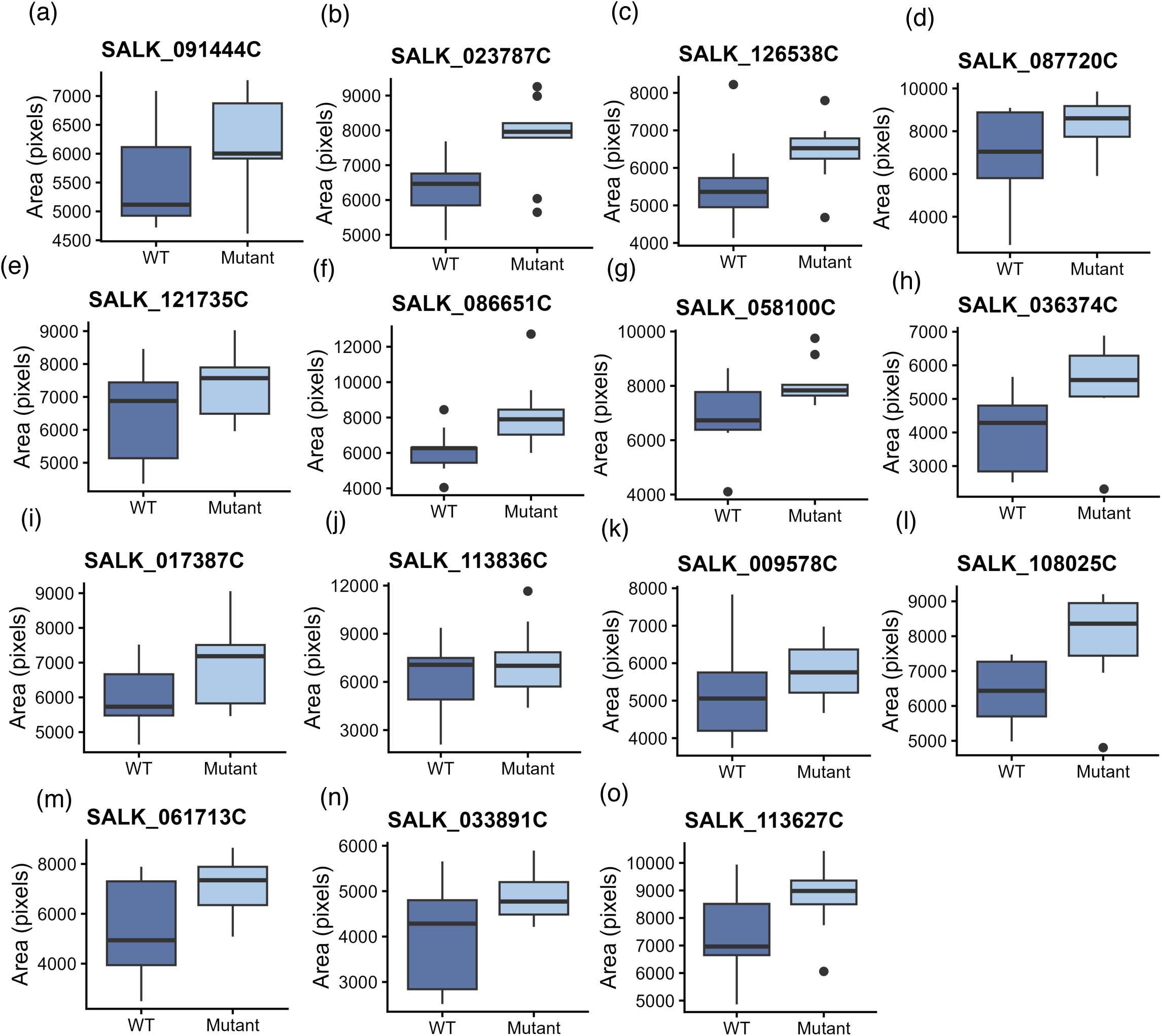
Enhanced callus growth in 15 T-DNA insertion lines. Boxplots show callus area (pixels) for 15 mutant lines with significantly larger calli than their respective plate-matched Col-0 controls (nominal P < 0.05). Each panel displays the SALK identifier of the mutant and its corresponding Col-0 control. The affected genes are (a) AT1G17030, (b) AT5G22000, (c) AT3G03773, (d) AT4G28220, (e) AT1G13960, (f) AT1G12160, (g) AT2G18280, (h) AT4G28260, (i) AT1G79120, (j) AT1G75540, (k) AT3G10500, (l) AT5G22090, (m) AT4G19860, (n) AT5G46460, and (o) AT4G13640. Panels are ordered to correspond to the same mutant lines shown in Figure 4.

### Candidate loci span diverse biological functions

Candidate loci identified in the screen encoded proteins associated with chromatin organization, transcriptional regulation, hormone signalling, metabolism, transport and cell wall-related processes. However, Gene Set Enrichment Analysis did not identify any category that remained significant after false discovery rate correction (Table S8,9; Figure S2). These results suggest that callus growth is influenced by diverse biological processes rather than a single dominant genetic pathway.

### Genetic determinants of reduced callus proliferation

To further resolve genetic contributors to callus proliferation, we examined the mutants displaying the strongest negative growth phenotypes. Among the 50 lines with the smallest calli (Table S4), 45 also exhibited reduced germination rates (1–3 out of 9 seeds germinated). The most severe phenotypes corresponded to mutations affecting genes encoding a MA3 domain-containing protein (AT3G48390) involved in translational regulation, the DNAJ-containing protein THERMOSENSITIVE MALE STERILE 1 (AT3G08970), the protein kinase AtPERK9 (AT1G68690), the RNA polymerase II elongation factor MDA7.18 (AT5G56120), and a heavy metal transport/detoxification protein (AT1G56210). These genes encode proteins involved in translation, protein folding, intracellular transport and transcriptional regulation

In contrast, a subset of mutants showing strongly reduced callus proliferation did not display impaired germination, suggesting more specific effects on post-embryonic proliferative capacity. These included mutations in the copper transporter COPT2 (AT3G46900), an unannotated SNARE-domain protein (AT4G22850), an oligomeric Golgi complex subunit-like protein (AT4G24840), a rhomboid-related intramembrane serine protease (AT5G25640), and a Mo25 family protein (AT2G03410). These loci represent candidates affecting proliferative growth after germination.

### Genetic determinants of enhanced callus proliferation

To further characterize the strongest candidates, we examined the subset of 15 mutant lines displaying significantly increased growth relative to their plate-matched WT controls (Figure 3). T-DNA insertions were distributed across distinct loci, in 1000 upstream, 5’ UTRs, exons and introns (Table S7).

The genes identified were associated with four broad and partially overlapping biological processes: mitochondrial and organelle function (NDB1, AT1G79120, AT5G46460), growth and developmental regulation (EAR1, RHF2a, BBX21, WRKY4, HSP90 co-chaperone p23-2), stress and nutrient signalling (ANAC053, EAR1, WRKY4, PHL3/UNE16, AT4G28260), and cellular and metabolic functions (TLP2, LCAT-PLA, FMOGS-OX7). Several of these genes remain poorly characterized despite having predicted molecular functions, including AT1G17030, AT1G79120, AT4G28260 and AT5G46460.

We next examined the molecular consequences of the T-DNA insertions by measuring transcript abundance directly in callus tissue. The effects of the insertions varied substantially among the selected lines. Four lines, AT5G22000 (AtRHF2a), AT5G22090 (EAR1), AT4G28260 (acyl-UDP-N-acetylglucosamine O-acyltransferase) and AT3G10500 (ANAC053), corresponded to knockout alleles. Three alleles corresponding to WRKY4 (AT1G13960), a calcium-dependent phospholipase (AT4G19860) and a co-chaperone HSP90p23-2 (AT3G03773) were significantly down-regulated.

In contrast, six genes were significantly upregulated: *PHL3/UNE16 (*AT4G13640), *NDB1(AT4G28220*, *FMOGS-OX7 (*AT1G12160), Pentatricopeptide repeat (PPR) superfamily protein (AT5G46460) AT1G79120 and AT1G17030, with expression ranging from 1.38 – to 4.27-fold that of the wild type. Two lines, corresponding to TLP2 (AT2G18280) and BBX21 (AT1G75540), transcript abundance did not differ significantly from the wild type despite their increased callus-growth phenotype. Therefore, the phenotype of these lines cannot be directly associated with altered expression of the corresponding genes. The possibility that the phenotype results from additional mutations present in these lines, or from effects of the T-DNA insertion not reflected in transcript abundance, cannot be excluded. Consequently, these genes were not considered further in the functional interpretation of the growth phenotype.

Among the high-growth candidates, *NDB1* (AT4G28220), encoding an external mitochondrial NAD(P)H dehydrogenase, has previously been implicated in vegetative biomass accumulation, with loss-of-function mutants showing reduced biomass ^18^. The SALK_087720C line identified in our screen carries a T-DNA insertion in the 5′ region and showed a 4.27-fold increase in *NDB1* expression in callus (p < 0.001), together with increased callus growth. Thus, the increased-expression phenotype observed here is consistent with the previously reported reduction in biomass following loss of *NDB1* function and identifies *NDB1* as a candidate for callus growth enhancement through enhanced expression.

AT4G13640 encodes PHR1-LIKE 3, a GARP-family transcription factor associated with phosphate-responsive regulation. In the SALK_113627C line, the T-DNA insertion is located approximately 1 kb upstream of the coding sequence. Expression analysis showed a 1.35-fold increase in AT4G13640 transcript abundance relative to wild type (p = 0.003), indicating that the enhanced callus-growth phenotype in this line is associated with increased expression of PHL3.

Several of the remaining genes have previously described functions that provide biological context for their increased callus-growth phenotypes. EAR1 was identified as a knockout in our screen, consistent with previous observations that loss of EAR1 function can promote growth under drought conditions ^19^. RHF2a was also identified as a knockout and has a redundant function with RHF1a during Arabidopsis gametogenesis ^20^. The p23-2 line showed strongly reduced expression (0.27-fold relative to WT); loss of Atp23-2 has previously been associated with reduced root growth and impaired root meristem maintenance ^21^. ANAC053, which was identified here as a knockout, encodes a membrane-associated NAC transcription factor implicated in ROS production during drought-induced leaf senescence ^22^.

Taken together, these candidates illustrate the diverse genetic basis of callus proliferation and provide multiple entry points for the genetic improvement of plant cell cultures.

### Enhanced Callus Growth Does Not Predict Vegetative Growth

To determine whether mutations promoting callus proliferation also enhanced whole-plant growth, selected high-growth mutants were grown in soil under standard conditions and vegetative growth was evaluated 28 days after sowing.

High-growth mutants displayed diverse vegetative phenotypes. Some lines exhibited larger rosettes than WT (Figure 4a–g), whereas others were smaller (Figure 4h–k) or showed no obvious differences from WT (Figure 4l–o).

**Figure 4.**
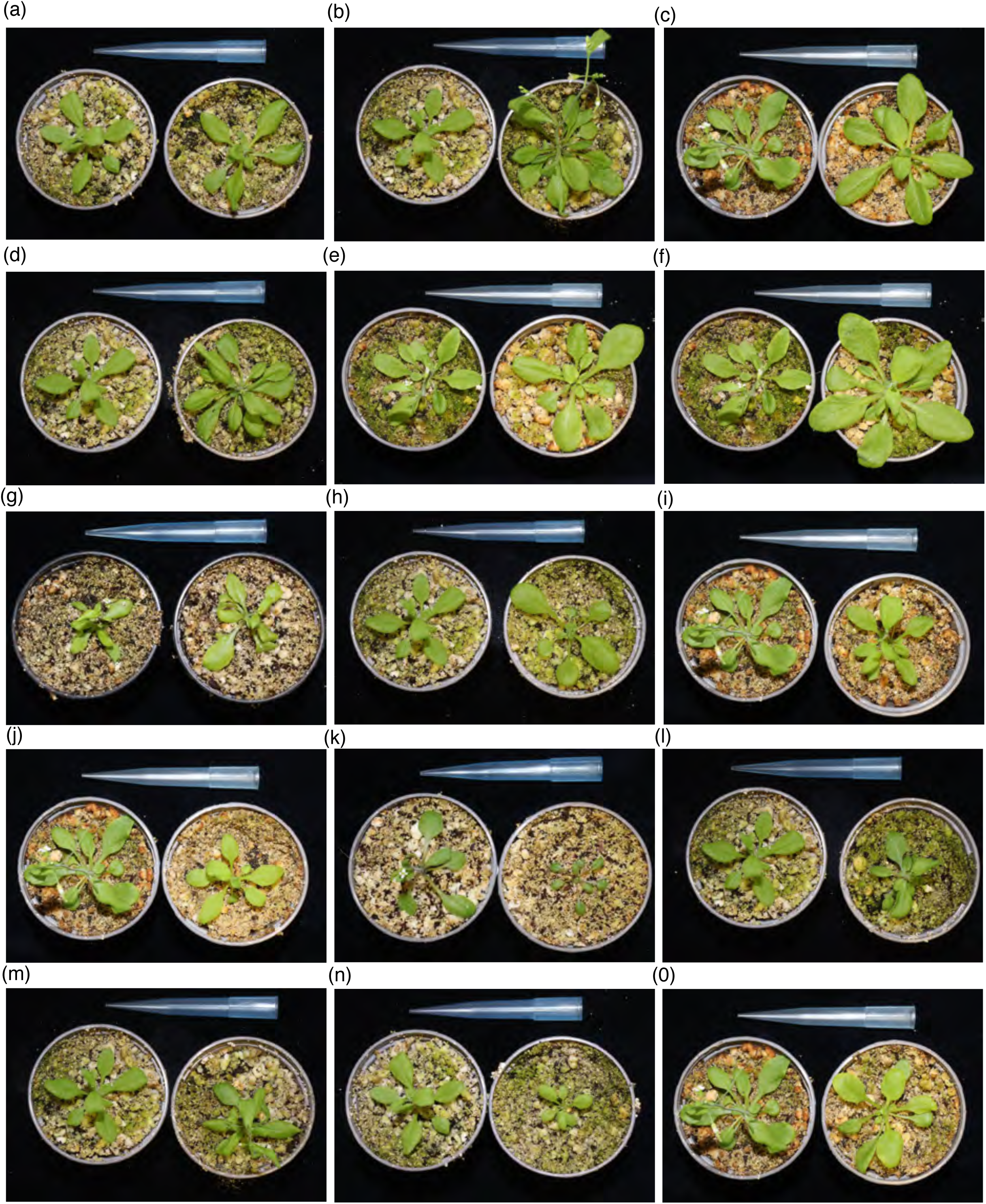
Enhanced Callus Growth Does Not Predict Vegetative Growth. Representative vegetative phenotypes of the 15 high-callus-growth mutant lines after two weeks of growth on plates followed by two weeks in soil. Col-0 is shown on the left of each panel and the corresponding mutant on the right. Images show one representative plant from ten plants analysed per genotype. (a) SALK_091444C (AT1G17030), (b) SALK_023787C (AT5G22000), (c) SALK_126538C (AT3G03773), (d) SALK_087720C (AT4G28220), (e) SALK_121735C (AT1G13960), (f) SALK_086651C (AT1G12160), (g) SALK_058100C (AT2G18280), (h) SALK_036374C (AT4G28260), (i) SALK_017387C (AT1G79120), (j) SALK_113836C (AT1G75540), (k) SALK_009578C (AT3G10500), (l) SALK_108025C (AT5G22090), (m) SALK_061713C (AT4G19860), (n) SALK_033891C (AT5G46460), (o) SALK_113627C (AT4G13640).

These observations demonstrate that enhanced callus proliferation is not necessarily accompanied by increased vegetative growth, supporting the conclusion that in vitro proliferative capacity can be genetically uncoupled from whole-plant biomass accumulation.

### High-Growth Callus Mutants Reveal Distinct Suspension Culture Phenotypes

To determine whether increased callus size on solid medium predicted improved suspension-culture performance, calli from five high-growth mutants were expanded on Petri dishes and equal amounts of biomass were used to inoculate liquid cultures. Growth trajectories were analysed using linear mixed-effects models fitted to log-transformed sedimented biomass measurements, allowing line-specific slopes to be interpreted as specific biomass accumulation rates (Figure 5).

**Figure 5.**
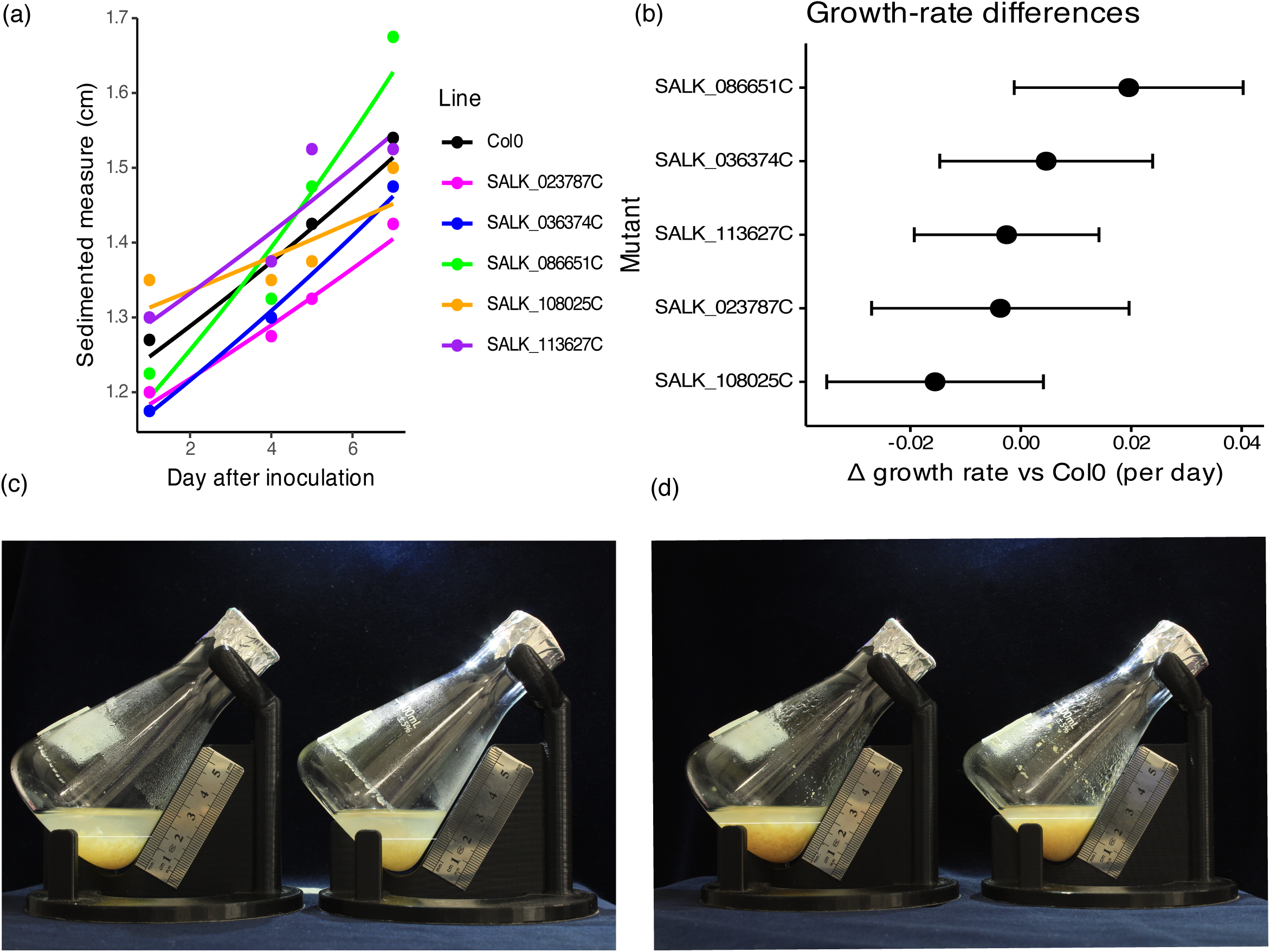
High-growth callus mutants reveal distinct suspension-culture phenotypes. (a) Growth trajectories of Col-0 and five high-callus-growth mutant lines in liquid culture: SALK_023787C (AT5G22000), SALK_036374C (AT4G28260), SALK_086651C (AT1G12160), SALK_108025C (AT5G22090), and SALK_113627C (AT4G13640). Growth trajectories were analysed using a linear mixed-effects model fitted to log-transformed packed cell volume (PCV) measurements. (b) Estimated growth rates for Col-0 and the five mutant lines, with confidence intervals derived from the model. Representative suspension cultures of Col-0 (left) and SALK_086651C (right) at (c) day 1 and (d) day 7 after inoculation. Images show representative flasks from eight Col-0 and four SALK_086651C biological replicates.

Two independent lines exhibited enhanced biomass accumulation relative to Col-0. After correction for initial inoculum volume, SALK_036374C and SALK_086651C accumulated approximately 1.27– and 2.47-fold the biomass of WT, respectively. These lines represent two different molecular outcomes of T-DNA insertion identified above: SALK_036374C is a knockout of AT4G28260, whereas SALK_086651C shows increased expression of FMOGS-OX7 (AT1G12160). Thus, improved suspension-culture growth was associated with both loss of gene function and increased gene expression.

During liquid-culture validation, maintenance of the cultures through successive passages revealed substantial differences in their capacity to sustain proliferation. Col-0 cultures showed a marked decline in proliferative capacity by the fifth passage and could no longer be maintained after the sixth passage. Several mutant lines lost growth capacity earlier, including SALK_108025C after three passages and SALK_086651C and SALK_113627C after four passages. In contrast, SALK_023787C and SALK_036374C remained vigorously proliferative through seven successive passages, when the experiment was terminated. These observations reveal an additional source of genetic variation among the selected lines and suggest that short-term biomass accumulation and the capacity to sustain proliferation across successive passages represent distinct, not necessarily coupled, components of suspension-culture performance.

Together, these results demonstrate that image-based phenotypic screening on solid medium can identify genetic variation affecting multiple traits relevant to suspension culture, including biomass accumulation and sustained proliferative capacity, providing a practical strategy for the genetic development of superior plant cell production lines.

## Discussion

Callus proliferation is central to regeneration, transformation, and plant cell–based manufacturing, yet optimization strategies have traditionally focused on media composition rather than genetic improvement. Here, we establish a scalable framework that treats callus growth as a quantitative genetic trait by integrating forward genetics with automated imaging and AI-assisted image analysis. This approach enabled the analysis of 27,648 calli and the identification of candidate loci modulating proliferative capacity, including rare mutations capable of enhancing growth.

Callus formation and proliferation are frequently regarded as predominantly hormone-driven processes ^23^. Although extensive work has focused on optimizing exogenous growth regulator combinations, genome-scale identification of genetic determinants controlling proliferative capacity has remained largely unexplored. By isolating callus directly from seeds under standardized conditions, we systematically quantified growth variation across 2,880 mutant lines and demonstrated substantial genetic control of proliferative capacity. The resulting phenotypic distribution was markedly asymmetric: while numerous mutations reduced callus size, only a small subset enhanced proliferation. Such a skewed distribution suggests that proliferative growth operates near a constrained optimum, where most perturbations disrupt functions required for growth, whereas rare mutations relieve regulatory constraints. Growth-enhancing alleles may therefore reflect the release of regulatory constraints rather than disruption of core biosynthetic processes, underscoring their potential value for biotechnological optimization. Together, the predominance of moderate-effect alleles and the limited number of loci remaining significant after multiple-testing correction suggest that callus growth is governed by a highly distributed genetic architecture rather than by a small number of major-effect regulators. The functional diversity of candidate loci, together with the absence of significantly enriched categories after multiple-testing correction, further supports a model in which callus growth is influenced by multiple biological processes rather than a single dominant pathway.

One initial working hypothesis was that enhanced callus growth would predominantly arise from mutations affecting core cell-cycle regulators. However, such genes were notably absent from the high-growth category. Instead, several loci associated with enhanced callus growth lacked functional annotation. This finding suggests that the genetic basis of enhanced proliferative capacity remains incompletely characterized and that forward genetic screening can uncover regulators that would not readily be predicted from current functional annotations.

A key outcome of this study is that enhanced callus growth does not universally translate into increased plant growth. Several mutants displaying larger calli exhibited reduced or unchanged vegetative growth under soil conditions, indicating that growth in dedifferentiated, hormone-rich environments can be genetically uncoupled from organ growth. These observations suggest that callus growth and whole-plant growth are at least partially independent traits, governed by overlapping but non-identical regulatory mechanisms.

The primary objective of this screen was to identify genetic determinants capable of improving growth in suspension systems relevant to bioreactor-based production. To assess the translational value of the primary screen, representative high-growth mutants were selected for liquid-culture validation. This validation was not intended as an exhaustive characterization of all candidate loci, but rather as a test of whether enhanced growth on solid medium could predict improved performance in liquid culture. Both SALK_086651C and SALK_036374C accumulated more biomass than Col-0 during the culture period. After correction for the initial inoculum volume, SALK_036374C and SALK_086651C exhibited approximately 1.27-fold and 2.47-fold greater net biomass gains than the wild type, respectively. Although these differences were measured over single seven-day culture cycles, they demonstrate that genetic variation identified through solid-medium screening can translate into altered biomass accumulation in suspension culture.

Even modest differences in growth rate can become consequential during repeated cultivation cycles as growth in liquid culture is exponential. Indeed, a 1.7-fold increase in daily growth rate has previously been associated with a reduction in bioreactor cultivation time from 21 to 10 days^24^. Viewed in this context, the 1.27-fold and 2.47-fold increases in PCV gain observed for SALK_036374C and SALK_086651C, respectively, highlight the potential value of genetically improving proliferative capacity in plant cell cultures. However, validation under bioreactor conditions will be required to determine the resulting effects on process productivity.

The diversity of candidate loci identified in the screen provides multiple routes for the genetic improvement of plant cell proliferation. While high-growth mutants identify loci whose disruption or altered expression can enhance proliferation, low-growth alleles may reveal positive regulators of proliferative capacity and therefore potential targets for overexpression or gene-dosage strategies. Beyond targeted engineering, these candidate loci may facilitate germplasm screening for naturally occurring allelic or expression variants associated with enhanced in vitro proliferation. The scale of the dataset therefore provides a resource for systematic prioritization of candidate loci and future combinatorial approaches to improving plant cell production systems.

Beyond short-term biomass accumulation, serial propagation revealed substantial genetic variation in the ability of cultures to sustain proliferation over successive passages. This trait is particularly relevant to long-term and continuous bioprocessing, where maintenance of a viable and proliferatively competent cell population is required over extended cultivation periods. Long-established plant cell lines such as tobacco BY-2, Arabidopsis T87 and our *Nicotiana benthamiana* LAB-3.1416JA line illustrate that plant cells can acquire or maintain proliferative capacity over very prolonged culture periods ^24–27^. However, exceptionally long-term proliferative capacity can only be established retrospectively, whereas sustained proliferation across serial passages provides a prospectively measurable phenotype for identifying candidate lines with extended proliferative potential. Importantly, our results indicate that this trait is not necessarily coupled to short-term biomass accumulation: SALK_086651C showed strongly enhanced biomass accumulation but lost proliferative capacity after four passages, whereas SALK_023787C and SALK_036374C retained vigorous proliferation through seven passages. Sustained proliferative capacity therefore emerges as a distinct and potentially important target for the genetic development of plant cell lines intended for extended and continuous production strategies, including perfusion-based cultivation.

## Concluding Remarks

Our study establishes callus proliferative capacity as a genetically tractable trait and provides a scalable framework for identifying genetic determinants relevant to plant cell biotechnology. Importantly, callus growth, whole-plant growth, short-term biomass accumulation, and sustained proliferation are not necessarily coupled, indicating that plant cell performance comprises distinct components that can potentially be optimized independently. Integrating genetic screening with automated phenotyping and suspension-culture validation therefore provides a route toward the development of plant cell lines tailored for in vitro production. Future work should determine whether these traits can be transferred across species, combined with high product yields, and maintained under bioreactor conditions.

## Experimental procedures

### Plant material and in vitro culture conditions

Arabidopsis thaliana T-DNA insertion lines from the SALK collection (NASC N27941; ABRC CS27941) were obtained from the Nottingham Arabidopsis Stock Centre (NASC) (Alonso et al., 2003). Columbia-0 (Col-0) was used as the wild-type control. For the proof-of-concept experiment, T-DNA lines affecting cell cycle-related genes were selected from the collection (Table S1). A total of 2,880 mutant lines were screened in the large-scale experiment (Table S2).

Seeds were surface-sterilized in 20% (v/v) commercial bleach containing 37 g L⁻¹ sodium hypochlorite and supplemented with 250 µL L⁻¹ Tween-20 for 20 min, rinsed four times with sterile distilled water, and stratified at 4 °C for 24 h.

For the proof-of-concept experiment, seeds were germinated directly on callus induction medium containing MS salts with B5 vitamins, 2,4-D and 6-BAP (Sello et al., 2017). Each plate was divided into 16 sectors containing 13 mutant lines and three Col-0 controls, with nine seeds sown per sector. Three independent plates were prepared (Figure S1).

For the large-scale screen, seeds were sown directly on callus induction medium in 12 × 12 cm square Petri dishes containing 70 mL of medium comprising MS salts with B5 vitamins, 30 g L⁻¹ sucrose, 10 mg L⁻¹ picloram, 0.2 mg L⁻¹ kinetin, and 3.5 g L⁻¹ Gelrite. Plates were divided into 16 equal sectors, with nine seeds of a single genotype sown in each sector. Each plate contained 15 mutant lines and one Col-0 control, with Col-0 consistently positioned in the upper-left sector. Plates were cultured at 23 °C under continuous light for 28 days before imaging.

### Image preprocessing and plate segmentation

Due to the high density of individuals per plate (144 seeds), two recurrent challenges were observed: (i) overlapping calli and (ii) growth extending beyond predefined grid sectors. To standardize image geometry and enable subsequent detection and spatial assignment of individual calli, a preprocessing pipeline based on classical computer vision methods was implemented prior to deep learning–based analysis.

Preprocessing steps included grayscale conversion, adaptive inverse thresholding, contour detection, and morphological filtering. The largest detected contour corresponding to the Petri dish was identified, and the image was cropped and geometrically normalized to ensure consistent scaling across plates. Each plate was then subdivided into 16 predefined spatial sectors corresponding to genotype positions.

This procedure enabled robust extraction of the plate region and established a standardized spatial framework for subsequent assignment of detected calli to genotype positions.

### Deep learning–based detection of calli and seeds

Object detection was performed using the YOLO11 architecture ^28^ trained to identify two classes: (i) calli and (ii) non-germinated seeds. A ground-truth dataset comprising 1,230 manually annotated calli along with associated non-germinated seed annotations from 26 Petri dish images was generated using the Labelme open-source tool (Wada et al., 2021). To avoid data leakage, the dataset was partitioned at the image level into training (18 images, n=851 calli), validation (5 images, n=254 calli), and testing (3 images, n=125 calli) subsets.

Multiple YOLO11 configurations were evaluated. All models demonstrated high generalization performance, with precision and recall values ≥0.96 and mAP_50_ values up to 0.994 on the test set. The YOLO11n configuration was selected for deployment due to its strong performance and computational efficiency.

Following bounding box detection, individual callus regions of interest were segmented using HSV color-space thresholding and contour extraction to generate precise pixel-level masks. The area was computed directly from the number of pixels contained within each mask. Because acquisition geometry and imaging parameters were fixed across experiments, pixel area was used as a consistent proxy for callus size.

### Growth quantification and data aggregation

Detected objects were assigned to genotypes based on their spatial position within each of the 16 predefined sectors. Non-germinated seeds were recorded for germination rate estimation but excluded from callus area calculations.

For each genotype on each plate, pixel areas of all detected calli (up to nine per genotype) were exported automatically to structured.xlsx files. Mean callus area for each genotype was calculated for downstream statistical analysis.

### Gene set enrichment analysis

We used the Webgestalt interface to identify enriched pathways using default settings ^29^.

### Vegetative growth assessment

For vegetative growth assessment, seedlings were grown vertically on plates at 23 °C under continuous light for 14 days and then transplanted to soil (perlite:vermiculite:substrate, 1:1:0.5). Plants were acclimated under a covered tray for 5–7 days and then grown in a 12 h light/12 h dark photoperiod (23 °C day/18 °C night). Plants were watered twice per week with ATM solution ^30^. Images were acquired at 28 days after stratification, corresponding to 14 days post-transplantation.

### Liquid culture analysis

Calli from selected high-growth lines were propagated on solid medium, mechanically disrupted, and used to inoculate liquid cultures. For each line, 4 g fresh callus biomass was used to inoculate 16 mL fresh medium in 100 mL Erlenmeyer flasks. Cultures were maintained at 23 °C under light conditions with shaking at 120 rpm. Biomass accumulation was monitored using a custom 3D-printed sedimentation device (HD-PLA) designed in SolidWorks and produced from an.STL file (Supplementary Material S1).

Packed cell volume (PCV) was measured daily by allowing cultures to sediment for 10 min. Cultures were subsequently maintained by serial subculture, transferring growing cultures into fresh medium under the same conditions. Successive subcultures were recorded as passages (P1, P2, etc.). When a culture lost proliferative capacity, the corresponding line was re-established in liquid culture from callus propagated on solid medium when additional biological replicates were required.

Growth trajectories were analysed using linear mixed-effects models fitted to log-transformed sedimentation measurements: log(PCV) ∼ Day × Line, with replicate as a random effect using R 4.3.2. Estimated slope differences between each mutant and Col-0 were obtained using emtrends (emmeans), and contrasts were computed using a treatment-versus-control comparison. Overall growth was calculated by subtracting the initial volume from the final volume for each sample.

### Gene expression analysis of T-DNA mutant lines

Leaf samples were collected from soil-grown plants of each line, callus samples were processed directly from in vitro cultures. Total RNA was extracted using the NucleoSpin ® RNA kit according to the manufacturer’s protocol. For the conversion to cDNA, we used the Maxima™ H Minus cDNA Synthesis Master Mix commercial kit containing DNases from ThermoFisher ©. Transcript presence or absence for candidate genes was assessed by endpoint RT-PCR in leaves. For selected lines showing expression, transcript levels were quantified by RT-qPCR using SYBR Green chemistry and normalized to SAND (AT2G28390) as the reference gene using callus tissue. Relative expression was calculated using the 2^-ΔΔCt method ^31^. We used at least three biological replicas and two technical per extraction. Primer sequences are provided in Supplementary Table S10.

## Acknowledgements

This work is funded by Proyecto PID2021-127933OB-C21; financed by MICIU/AEI/10.13039/501100011033 and FEDER to M.E.-C. and J.W.; and Catedra BIONET in Biotechnology, Grant DIN2020-011559, funded by MICIU/AEI/10.13039/501100011033 and by “European Union NextGenerationEU/PRTR”, to F.V.-N. We would like to thank Dr. Qikun Liu for comments on the manuscript.

## Author contributions

Conceptualization, F.V-N, J.W, J.K, P.J.N-L, D.P.B and M.E-C.; Methodology, F.V-N, J.W, J.K, P.J.N-L, D.P.B and M.E-C.; Investigation, F.V-N, A.E-V, J.W, J.K, P.J.N-L, D.P.B, D.B-M, E.S-F, and M.E-C.; Writing – Original Draft, F.V-N, J.W and M.E-C; Writing – Review and Editing, F.V-N, A.E-V, J.W, J.K, P.J.N-L, D.P.B, D.B-M, E.S-F, and M.E-C Funding Acquisition, F.V-N, J.W, and M.E-C.; Resources, J.W, J.K, P.J.N-L, D.P.B, and M.E-C; Supervision, J.W, J.K, P.J.N-L, and M.E-C.

## Use of AI

During the preparation of this work, the authors used ChatGPT (OpenAI) for English-language editing and to assist in the development and refinement of R scripts used for data analysis and figure preparation. The authors reviewed and verified all AI-assisted outputs and take full responsibility for the content of the manuscript.

All figures were prepared by the authors, using Inkspace and Biorender

## Supplemental information

Figure S1. Proof-of-concept analysis of selected cell-cycle mutants affecting callus development. (a) Plate layout used for the proof-of-concept experiment. Each plate contained nine seeds per sector, comprising 13 mutant lines and three Col-0 controls, for a total of 144 seeds per plate. Thirteen alleles representing eleven genes were analysed (Table S1). Col-0 sectors are indicated in blue, mutants producing smaller calli in orange, and the mutant producing larger calli in green. (b) T-DNA insertion positions in AT1G49620 (*AtKRP7*).

**Figure S2. Gene set enrichment analysis of lines with reduced and enhanced callus growth.**

**Supplementary material S1**. STL file to print the Erlenmeyer holder on a standard 3D printer-Available upon request

**Table S1.** Arabidopsis T-DNA lines affecting genes related to cell cycle used for proof of concept.

**Table S2**. Arabidopsis T-DNA lines tested from NASC collection.

**Table S3.** Arabidopsis T-DNA lines showing low or null germination rates.

**Table S4.** Arabidopsis T-DNA lines showing decreased growth compared to the WT average.

**Table S5.** Arabidopsis T-DNA lines showing increased growth compared to the WT average.

**Table S6**. Arabidopsis T-DNA lines showing significant smaller calli area than their respective plate control (WT).

**Table S7**. Predicted T-DNA insertion sites, primers, and expression analysis of high-callus-growth mutant lines.

**Table S8.** Results of Gene Enrichment Analysis for the genes affected by T-DNA insertions causing decreased growth compared to the WT average.

**Table S9.** Results of Gene Enrichment Analysis for the genes affected by T-DNA insertions causing increased growth compared to the WT average.

## Data availability statement

The data that supports the findings of this study are available in supporting information of this article.

## Declaration of generative AI and AI-assisted technologies in the manuscript preparation process

During the preparation of this work, the author(s) used CHATGPT 5.1 for English polishing and improve R-scripts for figures. The author(s) reviewed and edited the output as needed and take full responsibility for the content of the published article.

