## Supplemental table and figures for "Large-scale forward genetics reveals genetically separable components of plant cell proliferative capacity"

Supplementary Table S1. Arabidopsis T-DNA lines affecting genes related to cell cycle used for proof of concept.

| **Number** | **Mutant Line** | **Affected gene** | **Gene description** |
| --- | --- | --- | --- |
| 1 | SIM-3 | AT5G04470 | SIAMESE |
| 2 | SALK_029546C | AT5G64960 | CYCLIN DEPENDENT KINASE C2 |
| 3 | SALK_089717C | AT3G24810 | KIP-RELATED PROTEIN 5 |
| 4 | SALK_122748C | AT1G27450 | ADENINE PHOSPHORYL-BOSYL TRANSFERASE 1 |
| 5 | SALK_081956C | AT5G21280 | ATR7 |
| 6 | SALK_149280C | AT5G64960 | CYCLIN DEPENDENT KINASE C2 |
| 7 | SALK_026391C | AT2G23430 | KIP-RELATED PROTEIN 1 |
| 8 | SALK_090154C | AT5G39610 | ARABIDOPSIS NAC DOMAIN CONTAINING PROTEIN 92 |
| 9 | SALK_047329C | AT1G27450 | ARABIDOPSIS ADENINE PHOSPHORIBOSYLTRANSFERASE 1 |
| 10 | SALK_053533C | AT3G24810 | KIP-RELATED PROTEIN 5 |
| 11 | SAIL_694_C04 | AT5G39610 | NAC DOMAIN CONTAINING PROTEIN 2/6 |
| 12 | SAIL_343_H12 | AT1G49620 | KIP-RELATED PROTEIN 7 |
| 13 | SALKSEQ_123548.1 | AT1G49620 | KIP-RELATED PROTEIN 7 |

(a)

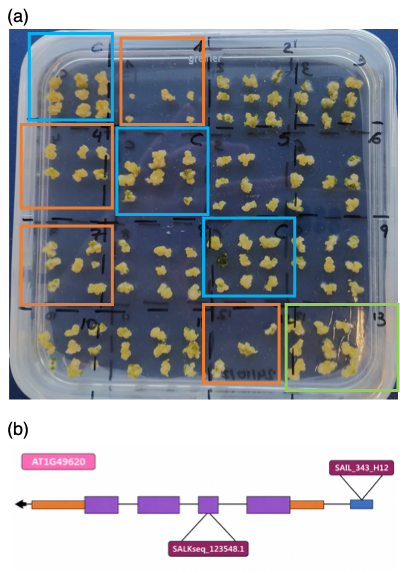

**Figure S1** (a) Plate showing differing growth rates compared to Col0 (C) on plates. Line 13 corresponds to KRP7 insertion SALKseq_123548.1 corresponding to a knockout.

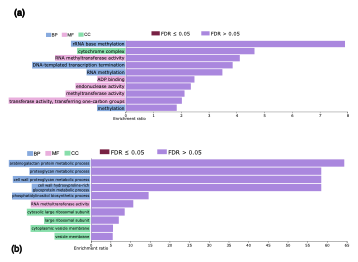

**Figure S2.** Gene set enrichment analysis of genes that upon mutation cause (a) smaller and (b) larger calli
